# Differential sensing with imprinted and non-imprinted organic electrochemical transistors reduces calibration drift in wearable sweat cortisol sensors

**DOI:** 10.64898/2026.10.31.763347

**Authors:** Lama I. Almofeez, Mohammad Shafiqul Islam, Thilina S. Ambagahawaththa, Yasser Khan

## Abstract

**Summary:** The output of a wearable cortisol sensor drifts over days for reasons unrelated to cortisol, so a calibration made on the first day soon stops being valid. In our differential organic electrochemical transistor, a PEDOT:PSS transistor coated with a molecularly imprinted polymer (MIP) for cortisol is paired with one coated with a non-imprinted polymer (NIP) of the same composition, and the NIP response is subtracted. Over four calibrations in artificial sweat across 14 days, the response of the MIP alone shifted by 15% to 22% of its range on day 1, whereas the differential output shifted by 1% to 10%, and the variation between days at 1 to 10 nM fell by 65% to 70% on average for individual pairs. In sweat from one participant, read against the calibration from day 1, both outputs tracked the direction of visible changes in salivary cortisol, while the MIP alone diverged from the differential output, consistent with the laboratory drift.

**The bigger picture:** Cortisol rises and falls each day and within tens of minutes of stress, and a sensor worn on the skin could follow these changes in a way that occasional blood or saliva samples cannot. To be useful, such a sensor has to stay accurate for days without recalibration, yet the soft polymers that make wearable chemical sensors sensitive also change slowly in contact with sweat. This work shows that a simple change in fabrication, placing a matched reference transistor beside the sensing transistor and subtracting the two signals, removes most of this slow change at low cortisol concentrations over two weeks. The reference is the non-imprinted polymer that sensors based on imprinted polymers already require as a control, so the approach adds no new materials and could carry over to imprinted sensors for other small molecules in sweat. Studies in more people with continuous wear are the next step toward use over several days.

## Introduction

Cortisol is the body’s main stress hormone and one of its most dynamic ^1^. Its level follows a daily cycle, rising sharply over the first 30 to 45 min after waking and falling through the rest of the day ^2^, and after an acute psychological stressor it peaks 21 to 40 min after onset ^3^. Disorders of the stress system appear as changes in its basal activity and in its responsiveness ^4,5^, so isolated blood or saliva samples capture only part of what clinicians and researchers want to know, and single measurements cannot distinguish acute from chronic stress ^1,6^. Sweat offers a way to follow cortisol repeatedly and without needles: it can be collected at the skin surface, stimulated on demand by iontophoresis, and analyzed by sensors worn on the body ^7,8,9^ (Figure 1B). In the past several years, wearable cortisol sensors have been built from molecularly imprinted organic electrochemical transistors (OECTs) ^10^, graphene immunosensors ^11^, and aptamer field-effect transistors ^12^, and multiplexed microfluidic patches now track cortisol together with epinephrine and norepinephrine ^6^. Wearable stress sensors are also being designed around where and how they are worn ^13^.

**Figure 1.**
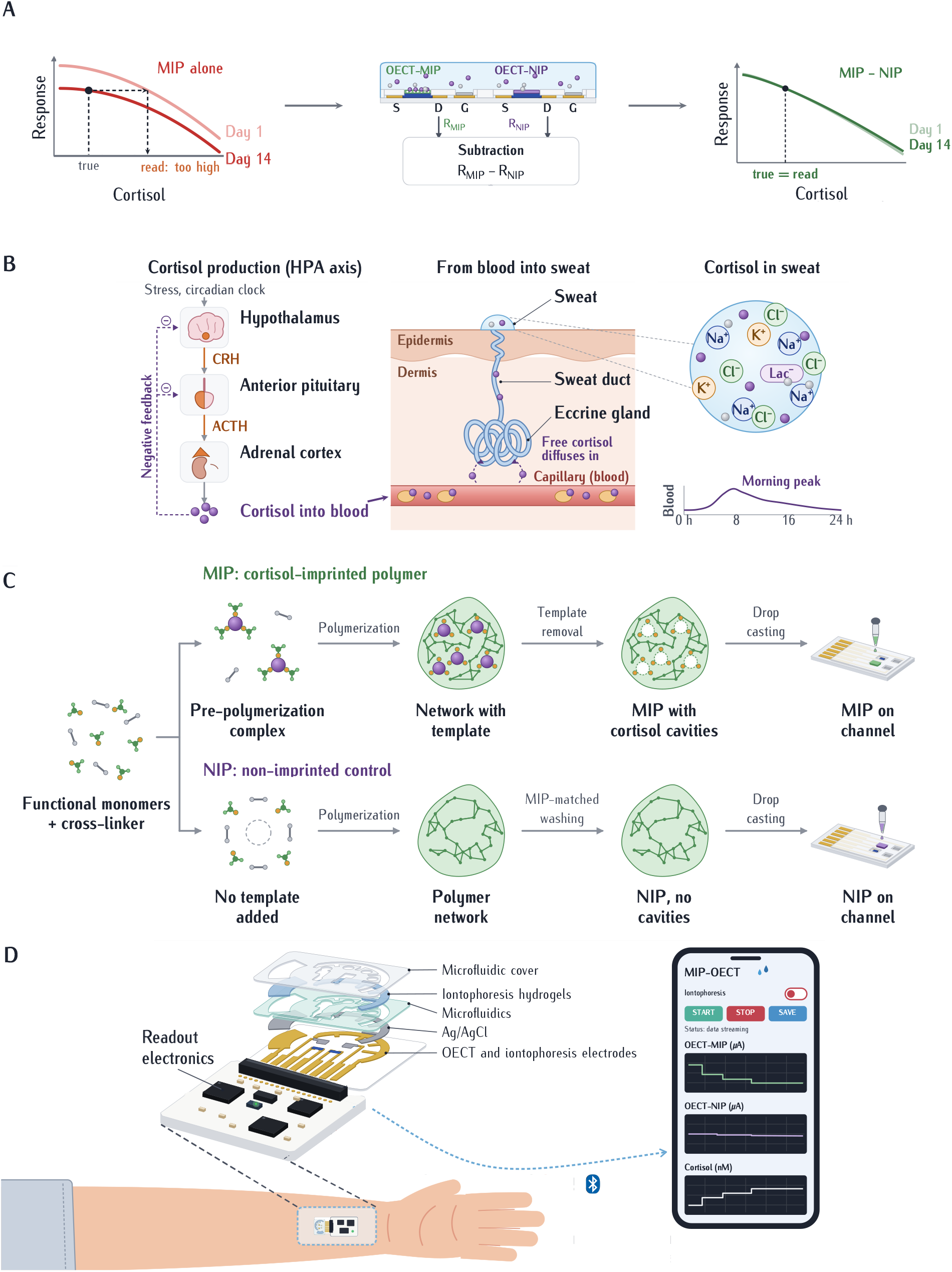
Concept of the differential MIP/NIP OECT. (**A**) Principle of the correction. Left, drift lowers the response of the MIP alone, so a reading taken against its day-1 calibration overestimates cortisol. Middle, the MIP- and NIP-coated OECTs share one sweat sample, and the NIP response is subtracted from the MIP response. Right, the MIP *−* NIP difference stays close to its day-1 calibration. (**B**) Cortisol released through the hypothalamic–pituitary–adrenal (HPA) axis diffuses from the blood into eccrine sweat. (**C**) Synthesis of the molecularly imprinted polymer for cortisol (MIP) and the non-imprinted control (NIP). (**D**) Wearable system: exploded view of the sensor patch, with the MIP- and NIP-coated OECTs and the iontophoresis electrodes on a flexible substrate, connected to wireless readout electronics worn on the forearm.

What these devices have not yet shown is that a sensor calibrated once can still be trusted days later. The obstacle is drift, a slow change in sensor output that is unrelated to the analyte. Reviews of wearable sweat sensing identify signal drift ^14^, biofouling ^14,15^, and variable sweat secretion ^9,14,15^ as obstacles to the sustained use of chemical sensors worn on the skin. Over several days the causes accumulate: the sensing layer swells, ages, and collects nonspecific deposits ^16^, and the composition of sweat varies over time ^15^. For cortisol, this problem is especially hard to work around. Drift and the hormone’s own daily rhythm unfold on the same timescale of hours to days, so drift cannot be filtered out the way fast electrical noise can. A sensor whose response slowly drifts reports a change in cortisol that never happened, and nothing in the signal itself reveals the error.

OECTs are well suited to this setting ^17,18^. Ions from the electrolyte enter the whole volume of the polymer channel, so small changes at the sensing interface become large changes in drain current, with transconductances in the millisiemens range at gate voltages below 1 V^17,19^. The same volumetric coupling that makes OECTs sensitive also makes them prone to drift. Poly(3,4-ethylenedioxythiophene):poly(styrene sulfonate) (PEDOT:PSS), the channel material used here, takes up water and ions and swells in contact with an electrolyte, and changes in its hydration, ion content, and structure alter the device response over time regardless of the target molecule ^16,20^. On the skin, where the electrolyte is sweat that arrives intermittently and varies in composition, these changes are difficult to prevent through materials design alone.

Molecularly imprinted polymers (MIPs) give OECTs chemical selectivity without biological reagents. Polymerizing a functional monomer and crosslinker around a template molecule, then removing the template, leaves cavities that match the target in size and shape. The resulting materials are synthetic, inexpensive, and stable under conditions that degrade antibodies and enzymes ^21,22^. Parlak and colleagues placed a membrane imprinted with cortisol over a PEDOT:PSS channel to build a wearable OECT in which cortisol binding reduces ion transport into the channel, and thus the change in drain current induced by the gate, and used it to detect cortisol in sweat on the body ^10^. Imprinted recognition is never perfectly specific, and the standard way to measure its nonspecific component is a non-imprinted polymer (NIP), prepared identically but without the template ^10,21^. In most MIP sensors, the NIP serves as a separate control that shows that the response comes from imprinting; in the device of Parlak et al., the transistor coated with the NIP did not respond to cortisol ^10^.

Pairing a sensing element with a matched reference that experiences everything except the target is an old idea in sensing. An ion-sensitive field-effect transistor can be measured differentially against a reference transistor whose surface does not respond to the ion of interest, so that disturbances common to both, such as an unstable potential of the reference electrode, cancel ^23^. Wearable organicfield effect transistors have recently been made largely insensitive to bias stress, temperature, and strain by pairing a transistor whose extended gate carries a cortisol aptamer with one that carries a scrambled control aptamer, and used to follow sweat cortisol during a cold pressor test ^24^. Electrochemical aptamer sensors in blood correct their drift by taking the difference between two signals that drift together but respond differently to the target: either two interrogation frequencies that respond in opposite directions ^25^, or two redox reporters, only one of which responds to the target ^26^. Among imprinted sensors, mass-sensitive and optical devices have been run with a non-imprinted or blank reference channel alongside the imprinted one ^27,28^, the response of an NIP electrode has been subtracted to cancel the nonspecific adsorption of interferents ^29,30^, and two electrodes imprinted for different targets have served as each other’s reference ^31^; two of these studies noted that the subtraction also suppressed baseline drift during a measurement ^29,31^. These approaches correct effects that arise within a single measurement session, over minutes to hours, and none was tested for whether the subtraction keeps a single calibration valid over days of intermittent use (Table S1).

A transistor coated with the NIP is a natural reference for an OECT based on a MIP. It is prepared from the same reagents with only the template omitted, so it adds no new materials, and it shares the channel material, the polymer chemistry, and the sample, lacking only the imprinted cavities. Whether it actually drifts like its imprinted partner is not obvious, however. In the work of Parlak et al., the imprinted polymer was nanoporous whereas the non-imprinted polymer was markedly less porous ^10^, and in an imprinted photoelectrochemical sensor the NIP film had a smaller specific surface area than the MIP film^29^, so the two films could swell, age, and foul differently. To our knowledge, an OECT coated with a NIP has not been used as a continuously subtracted reference in a wearable sensor or evaluated against a single calibration over two weeks.

Here we test this idea directly. We fabricated pairs of PE-DOT:PSS OECTs with Ag/AgCl gates, coated the channel of one transistor in each pair with a MIP for cortisol and the other with the corresponding NIP, and used the difference between their normalized responses as the sensor output. Three device pairs were calibrated in artificial sweat from 1 to 500 nM on days 1, 4, 9, and 14, so that the drift of each transistor, and of their difference, could be measured over two weeks. On the same four days, sweat was collected from one participant during a cold pressor test, desk work, and stationary cycling and measured with a separate sensor pair. These readings were converted to concentration using only the calibration from day 1 and compared with salivary cortisol. We show that subtracting the NIP removes most of the drift between days at the low end of the range, and that in human sweat the sensor follows the direction of change in salivary cortisol, while the reading of the imprinted transistor alone moves away from the differential reading over two weeks in a way consistent with this drift.

## Results

### Sensor design and sensing mechanism

The sensor consists of two printed PEDOT:PSS OECTs operated side by side on a flexible strip and read by wireless electronics worn on the forearm (Figure 1D). The channel of the sensing transistor is covered with a poly(vinyl chloride) (PVC) membrane containing MIP particles imprinted with cortisol, and the channel of the reference transistor with an identical membrane containing non-imprinted particles, a membrane-on-channel architecture also used in ion-selective OECTs ^32^, for which an empirical model of the response has been reported ^33^. The two polymers were synthesized from the same formulation and washed in the same way, so that any difference between them arises from the imprinting step (Figures 1C and S1). This difference is visible in their structure: scanning electron microscopy (SEM) shows a porous surface on the MIP particles, whereas the NIP particles are considerably denser (Figure 2D), in agreement with the morphology reported by Parlak et al. ^10^.

**Figure 2.**
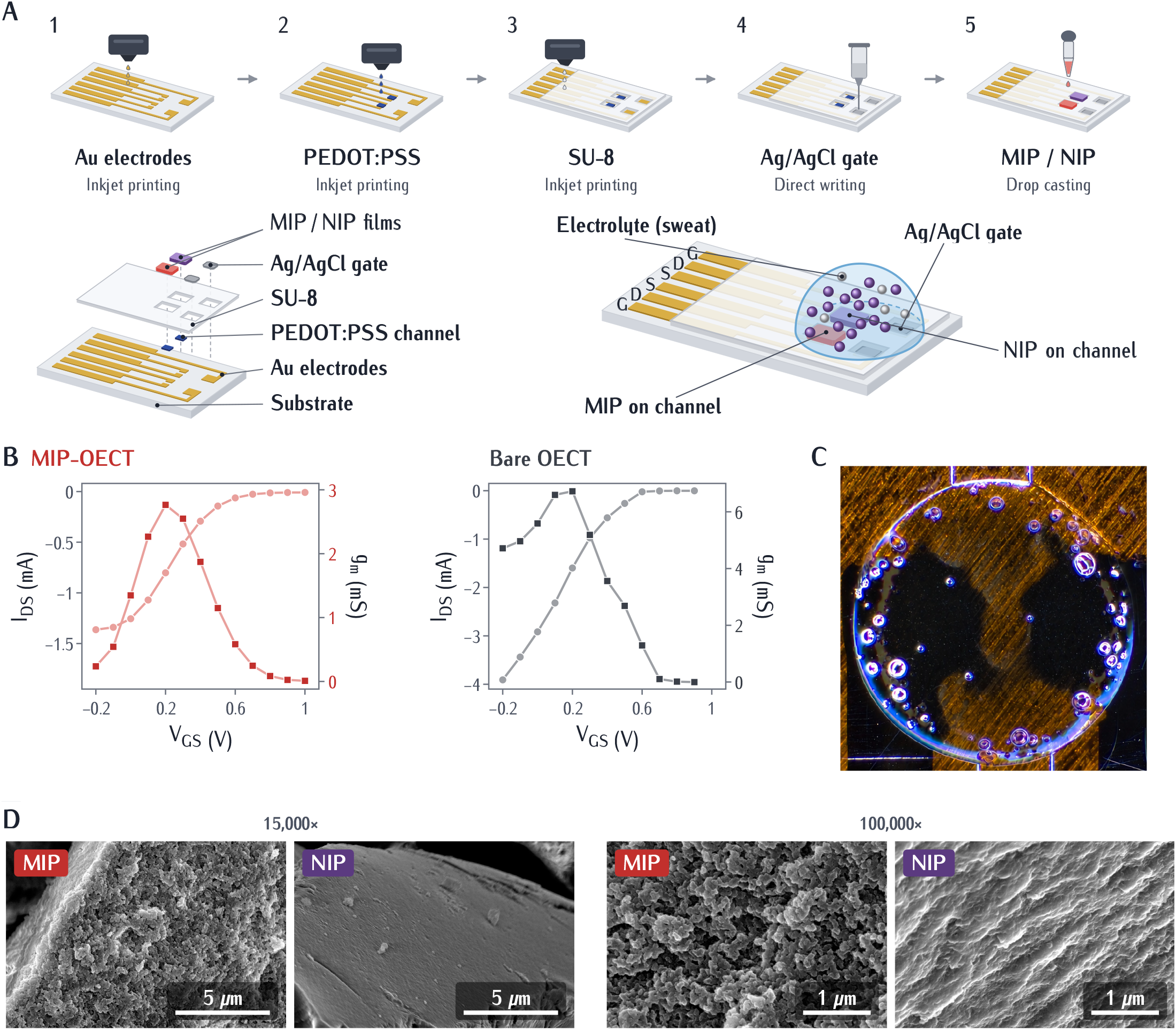
Fabrication and characterization of the MIP- and NIP-coated OECTs. (**A**) Printing sequence of one transistor pair, and exploded view of the layers and the device. (**B**) Transfer curves (*V*_DS_ = *−*0.5 V; circles, left axis) and transconductance (squares, right axis) of a MIP-coated and a bare OECT in artificial sweat containing 100 nM cortisol. Lines connect the measured points. (**C**) Optical image of the drop-cast MIP membrane. (**D**) Scanning electron microscopy (SEM) images of MIP and NIP particles at 15,000*×* (scale bars, 5 *µ*m) and 100,000*×* (scale bars, 1 *µ*m).

Each transistor is read by stepping the gate from 0 to 0.2 V at a fixed drain voltage and recording the drain current at each step. The response is the fractional change in drain current, *R* = |(*I*_DS_*−I*_0_)*/I*_0_|, where *I*_0_ is the current at *V*_GS_ = 0 V and *I*_DS_ the current at *V*_GS_ = 0.2 V^34^. When the gate is stepped, cations from the sweat enter the channel and dedope the PEDOT:PSS, lowering the current. Cortisol bound in the imprinted cavities is reported to reduce this ion transport through the membrane ^10^, so the gate dedopes the channel less and *R* decreases as cortisol increases. The NIP membrane lacks these cavities but shares the channel material, the membrane chemistry, and the sample, so changes that affect the two transistors equally should appear in both responses. The sensor output is therefore taken as the difference *R*_MIP*−*NIP_ = *R*_MIP_ *− R*_NIP_ (Figure 1A).

### Device fabrication and transistor characteristics

The OECTs were fully printed on a flexible substrate using the fabrication process for wearable OECTs established in our previous work ^34^ (Figure 2A). The MIP-coated device operated as a transistor, with a lower drain current at *V*_GS_ = 0 V (1.26 versus 2.92 mA) and a lower maximum transconductance (2.8 versus 6.7 mS) than the bare device, so that the transconductance relative to the baseline current was similar for the two (2.2 versus 2.3 V^*−*1^; Figures 2B and S2). The transconductance peak remained near *V*_GS_ = 0.2 V for both devices, so this gate voltage was used as the operating point for all sensing measurements. The NIP-coated OECT also operated as a transistor but was gated more weakly at this operating point: between *V*_GS_ = 0 and 0.2 V its current fell by 3.5%, compared with 36% for the MIP-coated device, and its transconductance peaked at 0.6 V rather than 0.2 V (Figures S3 and S4). This device came from the same fabrication batch as the calibrated pairs, and its weak gating at the operating point is consistent with the small NIP response measured on the wearable system (0.010 to 0.036; Figure 3C).

**Figure 3.**
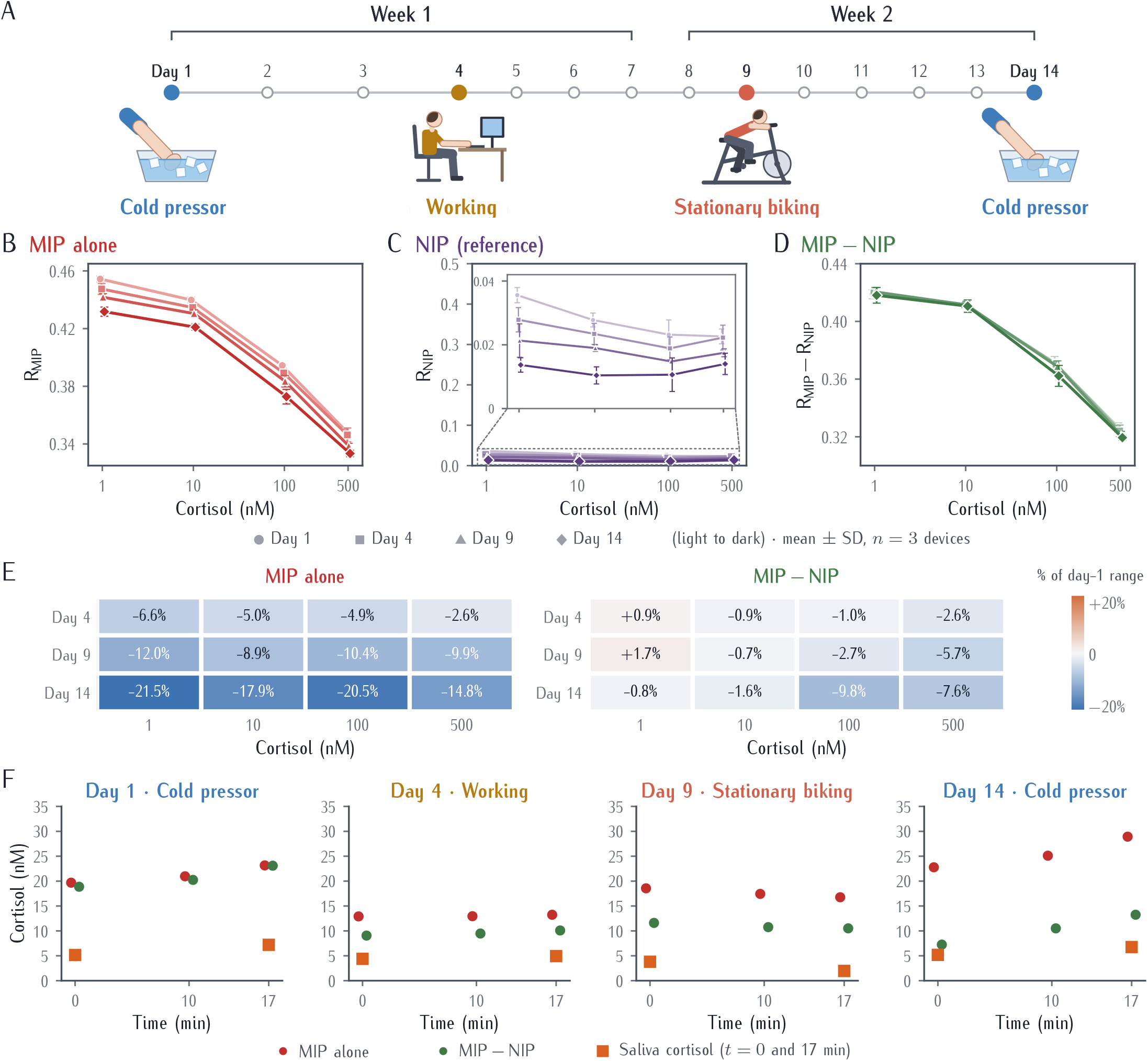
Calibration drift and measurements of human sweat over two weeks. (**A**) Study timeline. Three MIP/NIP pairs were calibrated in artificial sweat (1 to 500 nM) on days 1, 4, 9, and 14, and sweat was collected from one participant on the same days during a cold pressor test (days 1 and 14), desk work (day 4, labeled Working), and stationary cycling (day 9, labeled Stationary biking). (**B**) Response of the MIP alone. (**C**) Response of the NIP, plotted from 0 to 0.5 so that its size can be compared with the MIP response in **B**; inset, close-up of the boxed region. (**D**) Differential response, *R*_MIP_ *− R*_NIP_. In (**B**) to (**D**), points are mean *±* SD of *n* = 3 devices, with later days in darker colors. (**E**) Change in response from day 1, as a percentage of the day-1 calibration range, for the MIP alone and the differential output, each normalized to its own day-1 range. (**F**) Cortisol estimated from the MIP alone and from the differential output in sweat samples collected until *t* = 0, 10, and 17 min, using the day-1 calibration, with salivary cortisol at *t* = 0 and 17 min (squares).

### Response, selectivity, and operational stability

Measured with a source/measure unit (SMU), the response of the MIP-OECT decreased as cortisol increased from 100 pM to 1 *µ*M (Figures S5 and S6). Selectivity was tested against progesterone, a steroid structurally similar to cortisol, and epinephrine, which is released together with cortisol during stress and can therefore be present in sweat during the activities studied here. Cortisol was held at 100 nM while epinephrine or progesterone was added at 1 nM, 100 nM, or 1 *µ*M, and the normalized response *R* of each mixture was compared with the response to 100 nM cortisol alone (Figure S7A). All six mixtures read slightly below the cortisol-only response, by at most 1.2% (about 0.005), and across the three interferent concentrations the response varied by at most 0.002 for progesterone and 0.003 for epinephrine, with no consistent trend with concentration. By comparison, a 100-fold increase in cortisol, from 1 to 100 nM, decreased the response by 0.057 (Figure S6A). The largest deviation with an interferent present was therefore less than a tenth of this cortisol response, indicating that, in these SMU measurements, the MIP response is governed by cortisol rather than by these structurally or physiologically related molecules. When the concentration was stepped up from 1 nM to 1 *µ*M and back down, the responses on the way down returned close to those on the way up (Figure S8).

The response was reproducible over 100 consecutive measurement cycles on the SMU (Figure S9). During 8 h of operation in artificial sweat containing 100 nM cortisol, with one measurement every 15 min on the wearable system, the MIP-OECT response remained within 0.6% of its initial value (between 0.3925 and 0.3946; Figure S10), corresponding to estimated concentrations of 99 to 107 nM when read against the day-1 calibration of the wearable system (described below). Over the same period, the baseline current *I*_0_ rose by about 25%, so normalizing each reading to its own baseline removed this slow change in channel current within the session.

### Calibration and drift over two weeks

Normalizing the drain current change to each device’s own baseline current reduces the effect of device-to-device differences in channel conductance, which arise from variations in the printed PEDOT:PSS layer ^34^. On day 1, the MIP response decreased from 0.454 *±* 0.002 at 1 nM to 0.349 *±* 0.001 at 500 nM (mean *±* SD, *n* = 3 devices; Figure 3B). The calibration curve was not linear on a logarithmic concentration axis: the response changed by about 0.015 per tenfold increase in cortisol between 1 and 10 nM, and by about 0.065 per tenfold increase between 100 and 500 nM. The NIP response was small, between 0.010 and 0.036 over all days and concentrations, or 2% to 8% of the MIP response (Figure 3C). On day 1, the MIP response changed about 8.1 times more than the NIP response over the full concentration range.

Repeating the calibration on days 4, 9, and 14 (Figure 3A) revealed substantial drift in the MIP alone (Figure 3B). The calibration curve shifted downward over the two weeks at every concentration: the response fell from 0.454 to 0.432 at 1 nM, from 0.440 to 0.421 at 10 nM, from 0.394 to 0.373 at 100 nM, and from 0.349 to 0.333 at 500 nM. Because the shift was similar at both ends of the range, the change in response from 1 to 500 nM, which sets the sensitivity of the calibration, decreased only slightly, from 0.105 on day 1 to 0.098 on day 14 (7%). The NIP drifted as well (Figure 3C). At 1 nM its response fell from 0.036 to 0.014, the same change as that of the MIP, and its slight decrease with increasing cortisol on day 1 had disappeared by day 14.

Subtracting the NIP from the MIP removed much of this drift (Figure 3D; individual devices in Figure S11). Expressed as a fraction of the day-1 calibration range, by day 14 the response had shifted by 15% to 22% for the MIP alone across 1 to 500 nM, but by 1% to 10% for the differential output; at 1 nM the shifts were *−*21.5% and *−*0.8% (Figure 3E). The spread between days, measured as the difference between the highest and lowest daily value at each concentration, fell from 0.023 to 0.0024 at 1 nM and from 0.019 to 0.0015 at 10 nM, reductions of 90% and 92%. At 100 and 500 nM, the spreads of the mean curves fell by 58% and 55%. On average, the calibration range of the differential output did not shrink (0.092 on day 1 and 0.099 on day 14), compared with a 7% decrease for the MIP alone, but it varied between pairs, from a 1% decrease to a 21% increase (Table S2). The same metrics calculated for each device pair, with paired statistics, are given in Note S1 and Table S2; per pair, the spread fell by 65%*±*15% at 1 nM, 70%*±*3% at 10 nM, 49%*±*12% at 100 nM, and 53%*±*24% at 500 nM. The correction was not complete: at 100 and 500 nM the differential response also fell, from 0.371 to 0.362 and from 0.326 to 0.319, whereas at 1 nM it was unchanged (0.419 and 0.418).

### Measurements of human sweat compared with salivary cortisol

To test whether the differential correction holds in human sweat, sweat was collected from one participant in four 17-min sessions on days 1, 4, 9, and 14 (Figure 3A). In each session, sweat was stimulated by pilocarpine iontophoresis and collected with three Macroduct collectors, removed at *t* = 0, 10, and 17 min, so that each sample represents several minutes of secretion before its collection time (Experimental procedures). Each sample was measured with the same sensor pair, the test pair, which was not one of the three calibrated pairs. The cold pressor test, a controlled acute stressor that raises salivary cortisol ^35,36,37^, was performed on days 1 and 14, so that the response to the same stimulus could be compared at the start and end of the study. Desk work on day 4 and stationary cycling on day 9 represented everyday cognitive and physical activity. In every session, the readings were converted to cortisol concentration using only the day-1 calibration, and saliva, the standard noninvasive sample for cortisol in stress research ^38^, was collected at the start and end of the session for comparison.

In the three sessions in which salivary cortisol changed visibly (days 1, 9, and 14), both outputs changed in the same direction as salivary cortisol between the start and the end of the session (Figure 3F). During the cold pressor sessions, salivary cortisol rose 1.4-fold on day 1 and 1.3-fold on day 14; the differential estimate rose 1.2- and 1.8-fold, and the MIP-alone estimate 1.2- and 1.3-fold. On day 9, salivary cortisol fell about twofold and both estimates fell by about 10%. On day 4, salivary cortisol changed little (1.1-fold), and the change in the sensor responses (0.0004 to 0.0005) was at the level of the readout noise. The absolute estimates were higher than salivary cortisol. On day 1, when the calibration was fresh, the MIP alone read 19.7, 21.0, and 23.2 nM and the differential output 18.9, 20.2, and 23.1 nM at *t* = 0, 10, and 17 min, well above the salivary values of 5.2 and 7.2 nM. Because this difference was present before any drift could occur, it does not arise from drift see Discussion). On later days the two outputs diverged. From the first to the third sample, the MIP-alone estimate went from 12.9 to 13.2 nM on day 4, from 18.6 to 16.8 nM on day 9, and from 22.8 to 28.9 nM on day 14, whereas the differential estimate went from 9.1 to 10.1, from 11.6 to 10.5, and from 7.2 to 13.2 nM. The MIP alone therefore read 1.3- to 1.4-fold higher than the differential output on day 4, 1.6-fold higher on day 9, and 2.2- to 3.2-fold higher on day 14, compared with 1.0-fold on day 1. Because the test pair was not one of the calibrated pairs, its estimates also carry the device-to-device variation of the calibration. The SD of the three calibrated pairs on day 1 corresponds to about 1.1-fold in concentration for the MIP alone and 1.0- to 1.4-fold for the differential output over the range of the sweat estimates, much smaller than the difference from saliva. At 10 to 100 nM, where the differential response changes by about 0.041 per tenfold change in cortisol, the readout noise (0.0004; Experimental procedures) limits the resolution to about 1.02-fold.

The growing difference between the MIP alone and the differential output is consistent with the drift measured in artificial sweat. To test this, we used the laboratory curves of each day in three steps: the differential response was converted to a concentration with that day’s differential curve; the MIP response expected at this concentration was taken from that day’s MIP curve; and this expected response was read against the day-1 MIP curve, as the measured MIP response was. From the first to the third sample, this predicted MIP-alone estimate went from 11.9 to 12.3 nM on day 4, from 18.0 to 16.4 nM on day 9, and from 21.8 to 31.1 nM on day 14, and the observed MIP-alone estimates differed from these predictions by 2% to 8% of the observed values. The divergence between the two outputs in human sweat is therefore consistent with the drift characterized in the laboratory. This is a consistency check rather than an independent validation, because the differential output of the test pair serves as the reference. Because saliva was sampled only twice per session, the comparison with saliva tests the start and end of each session, not the time course between them.

### Continuous measurement with the worn sensor

To show that the sensor can also be read continuously on the body, a second participant wore a new sensor with its iontophoresis electrodes and microfluidic layer (Figures 4A and 4D) during a separate cold pressor session. After iontophoresis and filling of the microfluidic reservoir, the session began with a 6-min baseline, followed by a 2-min cold pressor test and 20 min of recovery (Figure 4E). Readings were taken during the baseline (*t* = 0 to 4 min) and during the last 10 min of recovery (*t* = 18 to 26 min); the MIP and NIP were read in alternate minutes, so each output was updated every 2 min (Experimental procedures). The recovery readings fall 12 to 20 min after the onset of the stressor. Cortisol peaks 21 to 40 min after the onset of psychological stressors ^3^, and in one study salivary cortisol peaked 10 to 20 min after a cold pressor test with a longer immersion than used here ^37^, so these readings may precede or overlap the peak of any response. Both outputs read 14.5 to 14.7 nM during the baseline and 15.6 to 16.6 nM during the recovery, about 13% higher at *t* = 26 min. The NIP response changed by less than 0.0005 over the session, so the change was in the MIP response. The change corresponds to about 0.0025 in the response, similar in size to the range of 0.0021 over 8 h at constant cortisol (Figure S10). Because no saliva was collected in this session, the change cannot be attributed to cortisol. The session shows that the worn sensor can be read continuously on the body. The two outputs agreed closely, as expected for a new sensor that had not yet drifted.

**Figure 4.**
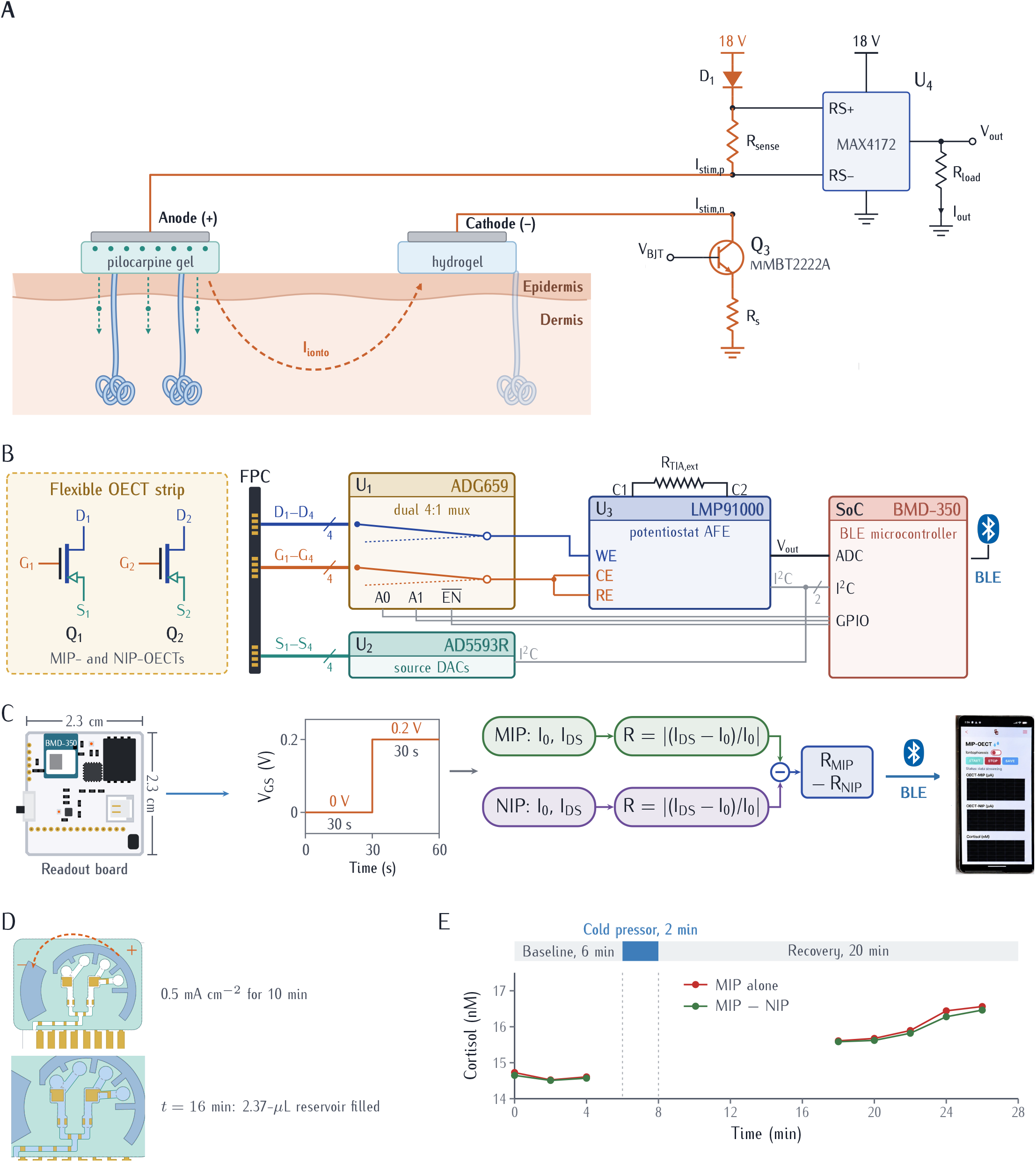
Wearable system. (**A**) Sweat stimulation by pilocarpine iontophoresis, and the constant-current iontophoresis circuit. (**B**) Multiplexed OECT readout circuit. (**C**) Readout board (illustration), gate-voltage sequence of one reading, and calculation of the differential output. (**D**) Sweat sampling on the patch: iontophoresis at 0.5 mA cm^*−*2^ for 10 min; the 2.37-*µ*L reservoir above the OECTs is filled by *t* = 16 min after the start of iontophoresis (channels shown in blue). (**E**) Cortisol estimated from the MIP alone and from the differential output with the day-1 calibration during a cold pressor session with the worn sensor: 6-min baseline, 2-min cold pressor, and 20-min recovery, read at *t* = 0 to 4 min and 18 to 26 min. The MIP and NIP were read in alternate minutes (MIP in the first minute, NIP in the second), giving one value of each output every 2 min.

## Discussion

A single calibration of a wearable chemical sensor is useful only for as long as the sensor’s response stays the same. Our results show that, for a PEDOT:PSS OECT with a MIP-containing PVC membrane, this period is short: within four days, the response of the MIP alone at 1 nM had shifted by 7% of its calibrated range, and by day 14 by a fifth. Pairing the MIP transistor with a matched NIP transistor and subtracting the two outputs removed most of this drift at low concentrations, and in human sweat the MIP alone moved away from the differential output over two weeks, consistent with this drift.

The drift of the MIP alone was a decrease in response at every concentration, by 15% to 22% of the day-1 range over two weeks, and the NIP drifted in the same direction. Over the two weeks, the NIP response changed by nearly the same amount as the MIP response at 1 to 10 nM and by about half as much at 100 to 500 nM, even though the NIP signal was only 2% to 8% of the MIP signal (Figures 3B and 3C). Cortisol bound in the imprinted cavities during earlier calibrations may remain there, because template bound in imprinted cavities can be hard to remove and occupied cavities are not available for rebinding ^39^; this lowers the response of the MIP and may have contributed to its fall, but it cannot explain a similar fall in the NIP, which has no cavities. The shared change was the part that the subtraction removed. At 1 to 10 nM, the two responses fell by about the same amount, rather than by the same percentage. The shared change therefore acted like an offset added to both outputs, not like a weakening of each membrane’s effect on its channel, which is why the subtraction removed it.

Drift in OECT biosensors arises from several processes that act at the same time ^16,40^, and the main ones in this device are shared by the two transistors of a pair. The first is the PEDOT:PSS channel. PE-DOT:PSS takes up water and swells in electrolytes, which changes the channel geometry, carrier mobility, and volumetric capacitance, and depletion-mode channels gradually dedope during operation ^16^; hydrophilic PSS, present in excess in printing inks, increases the swelling and the loss of current ^41,42^, and water retained in the film can cause irreversible changes ^40^. The membrane on top does not isolate the channel from these processes. Neutron reflectometry of a PEDOT:PSS solid contact beneath a plasticized PVC membrane showed that water passes through the membrane and is taken up by the hydrophilic PEDOT:PSS^43^, and ions must reach the channel for it to be gated at all. Because the devices were dried between calibration days, each calibration started from a new hydration of the channel. The second is the Ag/AgCl gate, a thin printed layer with no internal electrolyte. Such electrodes are depleted by dissolution of the thin AgCl layer, which exposes the conductor beneath and produces mixed potentials, and their potential follows the composition of the solution, including its chloride concentration ^44,45^; with an Ag/AgCl gate, the transfer curve of an OECT shifts with the ion concentration of the electrolyte ^46^. The third is the plasticized PVC mem brane. Plasticizer is released slowly from such membranes (no more than 2 ppm in solution after 12 h for bis(2-ethylhexyl) sebacate) ^47^ and can be transferred into the polymer beneath them^48^, and in ion-selective OECTs with a PVC membrane cast on the PEDOT:PSS channel, long-term stability depended on the mechanical stability of the membrane ^32^. The two transistors of each pair share the channel ink, the gate material and process, the membrane recipe, and the sample, so these changes are expected to be similar in both, which is the condition under which a matched reference cancels drift ^24^. Only processes specific to the imprinted cavities, such as the carry-over of cortisol between calibrations, act on the MIP alone and cannot be canceled by the NIP; they may account for part of the small drift that remained in the differential output.

We do not attribute the drift to a single cause. These processes act together and on similar timescales, few studies have examined the long-term stability of PEDOT:PSS devices ^49^, and the factors that limit the lifetime of miniaturized Ag/AgCl electrodes have been analyzed only sparsely ^45^. Separating their contributions would require experiments designed for that purpose. The differential design does not require knowing the cause: it removes the part of the drift that the two transistors share, whatever its origin. Other two-transistor designs have also suppressed drift strongly. Subtraction against a scrambled-aptamer reference reduced drift-induced distortion by up to two orders of magnitude in diode-connected organic field-effect transistors ^24^, and a dual-gate OECT, in which two transistors are connected in series rather than subtracted, largely mitigated drift ^50^. In our device, a small drift remained in the differential output.

The NIP was not fully concentration independent. On day 1, its response decreased slightly with increasing cortisol (slope *−*0.0049*±* 0.0005 per decade, about one-eighth of the MIP slope; Table S3), and by day 14 this slope had flattened on average (0.0000 *±* 0.0022 per decade), although with three pairs the change was not significant (one-sample *t* test on the change, *p* = 0.072). This slope is of the same size as its day-to-day variation, so we do not interpret it. The MIP slope changed less (+0.0019 *±* 0.0015 per decade, *p* = 0.16), and the slope of the differential output did not change significantly (*−*0.0030 *±* 0.0035 per decade, *p* = 0.27).

The correction was strongest at 1 to 10 nM, where the NIP drifted by as much as the MIP, and weaker at 100 to 500 nM, where it drifted by about half as much. The low end of the range is where the differential design matters most in practice: salivary cortisol in this study was about 2 to 7 nM, and mass spectrometry of eccrine sweat from one volunteer placed cortisol in a similar range, about 0.7 to 8 nM^51^. It is also where the calibration curve is flattest, so small errors in the response translate into large errors in concentration. Between 1 and 10 nM, a tenfold increase in cortisol changed the differential response by only about 0.007, so an error of 0.002 in the response shifts the estimated concentration by about twofold. For this reason, even the small drift that remained in the differential output matters at the low end of the range.

These results differ from the usual role of the NIP in MIP sensors. In most work, including the MIP-OECT of Parlak et al., the NIP is a separate control used to show that the response comes from imprinting ^10^. Here the NIP transistor is measured alongside the MIP transistor and subtracted from it, which turns the nonspecific part of the response from a confounding factor into a correction. The approach requires no change in materials or chemistry beyond preparing the NIP alongside the MIP, which is already standard practice.

In summary, a matched imprinted/non-imprinted transistor pair pro vides a simple way, built into the fabrication of the sensor, to reduce calibration drift in wearable cortisol sensing. Because it relies only on the NIP that MIP sensors already require, the same strategy could be applied to other imprinted sensors for small molecules in sweat.

### Limitations of the study

Calibration statistics are based on three device pairs, so the reported spreads are rough estimates. The test pair was read with a calibration measured on other devices; device-to-device differences between the calibrated pairs correspond to up to about 1.4-fold in concentration over the range of the sweat estimates. The sweat study involved one participant and eight saliva samples, and in each session the third sweat sample was collected from the other arm, so site-to-site differences in sweat cortisol may contribute to the change between the start and the end of a session. Salivary cortisol served as a reference for the direction of change, not for the absolute sweat concentration, because the relation between the two fluids is not established. Sweat and salivary cortisol were correlated in one study (*r* = 0.78)^11^ but not in another that used week-long sweat patches ^52^. Reported sweat concentrations also differ by more than a hundredfold between studies, depending on how the sweat was collected and measured: an immunoassay of sweat collected after intense exercise gave about 20 to 400 nM^53^, mass spectrometry of heat-induced eccrine sweat from one volunteer gave about 0.7 to 8 nM^51^, and in a study that measured both fluids in the same participants, sweat collected during exercise contained less cortisol than saliva ^54^. Sweat cortisol may also depend on sweat rate and hydration. The offset from salivary cortisol on day 1, before any drift, may therefore reflect the difference between the two fluids, differences between real and artificial sweat, or a difference between the test pair and the calibrated pairs; the present data cannot separate these. Over the two weeks, the MIP-alone estimates rose relative to salivary cortisol while the differential estimates fell relative to it; because the relation between sweat and saliva can change between sessions and collection sites, these data cannot show which output is closer to the true sweat concentration. In addition, the second saliva sample was taken 15 min after the onset of the 2-min cold pressor. In one study with a longer immersion, salivary cortisol peaked 10 to 20 min after the test ^37^, and after psychological stressors cortisol peaks 21 to 40 min after onset ^3^, so with only two samples per session the timing of our sample relative to the salivary peak is uncertain. The sweat results therefore test the direction of change and the relative drift of the two outputs; they are consistent with the drift correction but do not establish the absolute accuracy of the sensor or validate it for clinical use. Finally, the drift was tested on sweat collected in four sessions over 14 days, with each sample averaging several minutes of secretion, rather than with a sensor worn continuously. Studies with more participants and continuous wear will be needed to establish how long a single calibration remains valid in daily use.

## Experimental procedures

### Resource availability

#### Lead contact

Requests for further information and resources should be directed to and will be fulfilled by the lead contact, Yasser Khan.

#### Materials availability

Details of the materials and devices are available from the corresponding author on request.

#### Data and code availability

All data supporting the findings of this study are included in the article and its supplemental information. Any additional information required to reanalyze the data reported in this paper is available from the lead contact upon request.

### Matched MIP and NIP synthesis and membrane preparation

MIPs for cortisol were synthesized by bulk free-radical polymerization using the formulation of Parlak et al., with a 1:6:30 molar ratio of template, functional monomer, and crosslinker ^10^, which was also used by Daniels et al. ^55^. Because cortisol dissolves poorly in dichloromethane (DCM), methanol was used to dissolve it, as in Daniels et al. ^55^; here, cortisol was dissolved in methanol before the other components were added. Cortisol (36.25 mg, 0.10 mmol) in 1.0 mL of methanol was combined with methacrylic acid (MAA; 51 *µ*L, 0.60 mmol) as the functional monomer, ethylene glycol dimethacrylate (EDMA; 0.57 mL, 3.0 mmol) as the crosslinker, and azobisisobutyronitrile (AIBN; 5.75 mg, 0.035 mmol) as the radical initiator in DCM, corresponding to a cortisol:MAA:EDMA:AIBN molar ratio of 1:6:30:0.35^55^. The mixture was sealed, purged with nitrogen for 10 min to remove dissolved oxygen, which inhibits radical polymerization, and polymerized under UV irradiation at 20°C for 24 h. Non-imprinted polymers were prepared in the same way without cortisol. Both polymers were ground into particles and washed identically: the particles were dispersed in acetic acid/methanol (8:2, v/v), stirred for 30 min, and collected by centrifugation at 1,000 rpm for 15 min. This cycle was repeated five times, and the particles were dried under vacuum overnight (Figure S1).

To form the sensing layers, the dried particles were embedded in a plasticized PVC membrane following Parlak et al. ^10^. High-molecular-weight PVC (50 mg), bis(2-ethylhexyl) sebacate (120 *µ*L), potassium tetrakis(4-chlorophenyl)borate (15 mg), and 20 mg of MIP or NIP particles were dissolved or dispersed in tetrahydrofuran (2 mL). The MIP and NIP membranes had identical compositions, and 0.4 *µ*L of each mixture was drop-cast onto the channel of the sensing or reference transistor, respectively, and dried for 24 h at room temperature (Figure 2C).

### Device fabrication

The OECTs were fully printed on a flexible substrate as described previously ^34^, with inkjet-printed gold electrodes and a PEDOT:PSS channel (channel length 400 *µ*m, width 500 *µ*m; PEDOT:PSS formulation, thickness, annealing, and gate geometry as in Islam et al. ^34^), an inkjet-printed SU-8 layer passivating the metal tracks, and a direct-written Ag/AgCl gate for each transistor. The SU-8 layer prevents electrochemical reactions and leakage currents at metal exposed to the electrolyte, leaving only the channel and the gate open. Ag/AgCl was used as a non-polarizable gate, so that the applied gate voltage acts on the channel instead of being lost at the gate– electrolyte interface ^17^. The MIP and NIP membranes were then drop-cast onto the channels as described above. The MIP and NIP transistors of each pair were printed on the same strip and were covered by the same drop of solution during calibration and by the same sweat sample.

### Wearable readout system

The OECT readout circuit is shown in Figure 4B. The two OECTs, *Q*_1_ and *Q*_2_ (p-type, depletion mode), are the devices under test. The configurable ADC/DAC AD5593R (*U*_2_), controlled by the system-on-chip (SoC) through the I^2^C bus, generates the source bias voltages, and the micropower programmable potentiostat analog front end LMP91000 (*U*_3_) sets the gate and drain bias voltages and converts the drain current to an output voltage *V*_out_; the current-measurement circuit is described in Note S2 and Figure S12. Because the two transistors are driven and measured through the same readout path, the gate-drive (CE/RE) and drain (WE) terminals of *U*_3_ are switched between them by the dual 4:1 analog multiplexer ADG659 (*U*_1_).

During a measurement, *U*_1_ is first disabled by driving its active-low EN pin high, isolating the transistors from the LMP91000 readout path. The source bias is then configured through *U*_2_, and the LMP91000 bias and transimpedance amplifier settings are configured through its I^2^C interface. The required transistor is then selected with the A0 and A1 address inputs of *U*_1_, and the selected signal paths are enabled by driving EN low. The analog output voltage *V*_out_ of *U*_3_ is sampled by the internal ADC of the SoC (BMD*−* 350). This sequence is repeated alternately for the two transistors. For each transistor, the firmware computes the normalized response *R* = |*I*_DS_ *− I*_0_|*/I*_0_ from the currents averaged over about 10 s near the end of the 0 V and 0.2 V phases, and then subtracts the NIP response from the MIP response to obtain *R*_MIP*−*NIP_ (Figure 4C). The subtraction is therefore always performed on normalized responses, never on raw currents. A mobile application, connected to the board by Bluetooth Low Energy, starts and stops iontophoresis, sets its current, and starts and stops measurements, during which it streams the normalized responses and their difference.

Because the iontophoresis electrodes lie close to the transistors, stimulation and readout are sequenced. Before stimulation, *U*_1_ is disabled, isolating both transistors from the readout circuit, so that stimulation current cannot pass through the skin into the readout path and return through the lower *−* resistance measurement circuitry. A boost converter then generates an 18-V supply, and the base voltage of the current-control transistor is set by a DAC output of *U*_2_. The iontophoretic current flows from the 18-V supply through the current-limiting diode (about 1 mA), the current-sense resistor, the anode, the skin, the cathode, the current-control transistor, and its emitter resistor to ground. At the end of stimulation, the base voltage is set to zero to stop the current, and the boost converter is then disabled.

### Electrical measurements and calibration

Device operation was first validated with a Keysight B2902A precision source/measure unit (SMU). Output characteristics were recorded for *V*_GS_ from *−*0.2 to 1.0 V and *V*_DS_ from 0 to *−*1.0 V, and transfer characteristics at *V*_DS_ = *−*0.5 V, for bare, MIP-coated, and NIP-coated devices in artificial sweat containing 100 nM cortisol (Figures 2B and S2–S4). The time response, initial calibration, selectivity, and reversibility measurements (Figures S5–S8) were also performed with the SMU: each device was biased at *V*_DS_ = *−*0.5 V, first at *V*_GS_ = 0 V for 120 s to obtain *I*_0_ and then at *V*_GS_ = 0.2 V for 120 s to obtain *I*_DS_. The calibrations on days 1, 4, 9, and 14 and all measurements of human sweat were performed with the wearable readout system using the same bias conditions with shorter phases of 30 s each, so that a full reading was completed within each 1-min measurement interval. In both cases, the response was calculated as

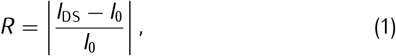

where *R* is shown as |Δ*I/I*_0_| on the axes of some supplemental figures. On the wearable system, each current was averaged over 318 samples (about 10 s at 32 samples per second) near the end of its phase, samples 584 to 901 for *I*_0_ and 1,483 to 1,800 for *I*_DS_ (Figure S13). The differential output is

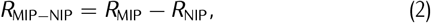

labeled MIP *−* NIP in figures. Because the response continues to change slowly during the gate pulse (Figure S5), absolute responses differ between the SMU and wearable protocols, so sweat readings were converted to con centration only with calibrations measured on the wearable system. The 100-cycle test (Figure S9) was performed with the SMU in artificial sweat containing 100 nM cortisol, with shorter phases of about 12 s.

Calibrations were performed in artificial sweat (BZ321, pH 6.3, Biochemazone) at 1, 10, 100, and 500 nM cortisol on three MIP and three NIP devices, forming three MIP/NIP pairs, on days 1, 4, 9, and 14. Concentrations were applied in increasing order, and each reading began immediately after the solution was applied. Bench measurements were performed at room temperature. On the wearable system, each solution drained through the microfluidic outlet before the next concentration was applied; in the benchtop SMU calibration, the previous drop was removed before the next was applied. For the reversibility test (Supplementary Fig. S8), the concentration was instead raised by adding cortisol to the drop on the device and lowered by diluting the drop with artificial sweat. No rinsing step was used between concentrations, to remain close to continuous operation on the body, in which sweat flows through the channel without rinsing. Between calibration days, devices were dried without washing and stored at room temperature in a closed, unsealed box. Selectivity was evaluated in artificial sweat containing 100 nM cortisol to which progesterone or epinephrine was added at 1 nM, 100 nM, or 1 *µ*M, keeping the solution volume equal for all measurements.

### Human participants

The study was approved by the University of Southern California Institutional Review Board, and both participants gave written informed consent before taking part. One volunteer, who was not an author, completed four sessions over 14 days: a cold pressor test on days 1 and 14, desk work (computer*−*based data processing) on day 4, and stationary cycling on day 9. The test pair was a separate MIP/NIP pair, not one of the three calibrated pairs, and the same pair was used in all four sessions. Its readings were converted to concentration using the mean day-1 calibration of the three calibrated pairs, measured on the wearable system in artificial sweat with known cortisol concentrations. All sessions were held between 12:00 and 2:00 p.m. to limit the influence of the daily cortisol rhythm.

Each session lasted 17 min and began with a 2-min resting baseline. During the cold pressor sessions, the participant immersed the right hand in ice water for 2 min, from *t* = 2 to *t* = 4 min, and the session continued until *t* = 17 min. During the desk-work session, the participant performed computer-based data processing and analysis from *t* = 2 to *t* = 17 min, and during the cycling session the participant cycled on a stationary bicycle at maximal self-selected effort over the same period to raise heart rate. Sweat samples were collected as described below. A second volunteer completed a separate cold pressor session with the worn sensor (Figure 4E): the participant rested for 6 min, immersed the right hand in ice water for 2 min (*t* = 6 to 8 min), and rested for a further 20 min; the sensor was read at *t* = 0 to 4 min and *t* = 18 to 26 min.

### Sweat stimulation and sampling

#### Collected sweat (Figure 3)

In the four sessions of the drift study, sweat was stimulated and collected with the Macroduct Sweat Collection System (Model 3700 Webster Sweat Inducer with Pilogel iontophoretic discs containing 0.5% pilocarpine nitrate; Wescor), which applies 1.5 mA for 5 min. Three samples were collected in each session, each from a separately stimulated site on the inner side of the arm. The first was stimulated from *t* = *−*20 to *−*15 min and collected until *t* = 0, the second was stimulated from *t* = *−*5 to 0 min and collected until *t* = 10 min, and the third was stimulated from *t* = 0 to 5 min and collected until *t* = 17 min, where *t* = 0 is the start of the session. Because sweat began to enter the collector about 3 to 5 min after stimulation, each sample represents between about 5 and 12 min of secretion before its collection time. The first two samples were collected from the left arm and the third from the right arm, above the part immersed in ice water during the cold pressor test. Each sample was applied to the same sensor pair immediately after collection and read on the wearable readout system in the same way as the calibration solutions; as in the calibrations, the sensor was not rinsed between samples.

#### Continuous measurement (Figure 4)

For the continuous measurement, sweat was stimulated by the wearable system. The iontophoretic electrodes (inkjet-printed Au with direct-written Ag/AgCl), the agarose anode hydrogel containing 5% (w/v) pilocarpine, the cross-linked poly(vinyl alcohol) cathode hydrogel, and the laser-patterned microfluidic layer were fabricated as described previously ^34^. The board supplies a constant iontophoretic current, set from the smartphone application, through a transistor current sink, limits it to a maximum of 1 mA with a series current-limiting diode, and monitors it in real time with a current-sense amplifier (see Wearable readout system). A current density of 0.5 mA cm^*−*2^over the 0.161-cm^2^ anode (about 80 *µ*A; cathode area 0.129 cm^2^) was applied for 10 min, delivering a total charge of about 48 mC, and iontophoresis ended before the first sensor reading. Stimulated sweat was guided to the transistor channels by the microfluidic layer. The MIP and NIP transistors were read in alternate minutes, so a differential value was obtained every 2 min. Although the two readings were 1 min apart, they sampled essentially the same sweat: the 2.37-*µ*L reservoir above the channels is filled by *t* = 16 min after the start of iontophoresis, so only about a tenth or less of this sweat is replaced within 1 min, whereas cortisol changes over tens of minutes after a stressor ^3^.

### Saliva collection and analysis

Saliva was collected at *t* = 0 and *t* = 17 min by placing a SalivaBio Oral Swab (item 5001.02, Salimetrics) under the tongue for 2 min. Swabs were transferred to Salimetrics Swab Storage Tubes (item 5001.05) and stored at *−*20°C until analysis. Salivary cortisol was measured with the Salimetrics Salivary Cortisol ELISA Kit (item 1-3002) according to the manufacturer’s protocol.

### Data analysis

Sweat cortisol was estimated by locating each normalized response on the mean day-1 calibration curve of the three calibrated pairs for the corresponding output (MIP alone or differential), using linear interpolation between neighboring calibration points on a logarithmic concentration axis. To predict the MIP-alone estimate from the differential output on days 4, 9, and 14, the differential response was converted to a concentration with the laboratory differential curve of the same day, the MIP response at this concentration was interpolated from the laboratory MIP curve of the same day, and this response was read against the day-1 MIP curve. The day-to-day spread at each concentration was calculated as the difference between the highest and lowest daily value, both from the mean curves and for each device pair. All sweat estimates (7.2 to 28.9 nM) fell within the calibrated range (1 to 500 nM), so no extrapolation was needed. Calibration data are reported as mean *±* SD of *n* = 3 devices.

#### Readout noise

One step of the readout ADC corresponds to about 0.002 in *R* for a single sample at the typical *I*_0_ of 2.0 to 2.6 mA, and each current is averaged over 318 samples. The readout noise, estimated as the SD of 33 repeated readings of the same solution over 8 h, was 0.0004 in *R* (Figure S10), about 12 to 95 times smaller than the NIP response of individual devices (0.005 to 0.038).

#### Statistical analysis

Drift metrics were also calculated separately for each device pair, pairing each MIP with the NIP of the same pair, and are reported as mean *±* SD across the three pairs. Calibration slopes were obtained for each device and day by linear regression of the response on log_10_ concentration. MIP-alone and differential metrics of the same pair were compared with two-sided paired *t* tests, and day-1 to day-14 slope changes were tested against zero with two-sided one-sample *t* tests.

## Acknowledgments

This work was supported by the National Institutes of Health (R61MH135407).

## Author contributions

L.I.A. and Y.K. conceived the project. L.I.A. designed the cortisol MIP- and NIP-OECTs, designed and performed the benchtop and on-body experiments, prepared and submitted the IRB application, managed the experimental logistics, and analyzed the data. M.S.I. helped with OECT printing. T.S.A. designed and programmed the printed circuit board. L.I.A. and T.S.A. wrote the manuscript. Y.K. supervised the project.

## Declaration of generative AI and AI-assisted technologies in the writing process

During the preparation of this work, the authors used Claude (Anthropic) to improve the clarity and grammar of the text. After using this tool, the authors reviewed and edited the content as needed and take full responsibility for the content of the publication.

## Supplemental information

Document S1. Figures S1–S13, Tables S1–S3, Notes S1 and S2, and supplemental references.

