## Supplemental Information for "Differential sensing with imprinted and non-imprinted organic electrochemical transistors reduces calibration drift in wearable sweat cortisol sensors"

### This file includes

Figures S1 to S13
Table S1 (differential and internal-reference strategies) and references for Table S1
Note S1 and Table S2 (per-pair drift metrics)
Table S3 (per-device calibration slopes)
Note S2 (transistor current-measurement circuit)

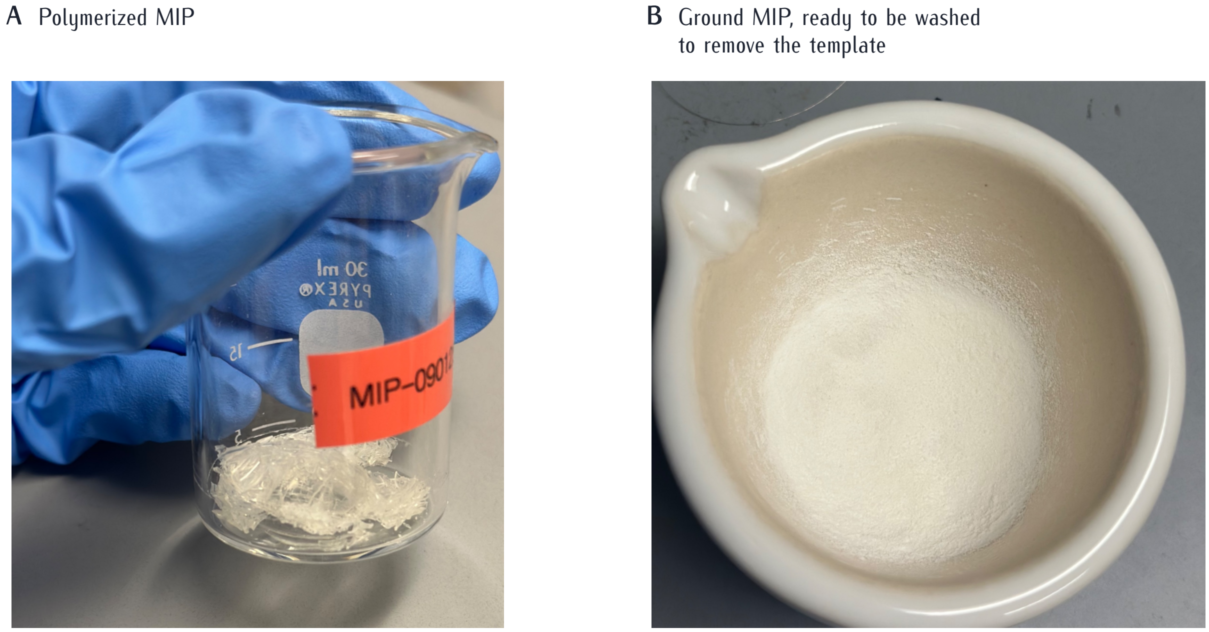


**Figure S1. Preparation of the MIP.** (**A**) MIP after 24 h of UV-initiated bulk polymerization. (**B**) Ground MIP particles before template removal in acetic acid/methanol (8:2, v/v). The NIP was prepared in the same way without cortisol. Photographs are unedited except for cropping.

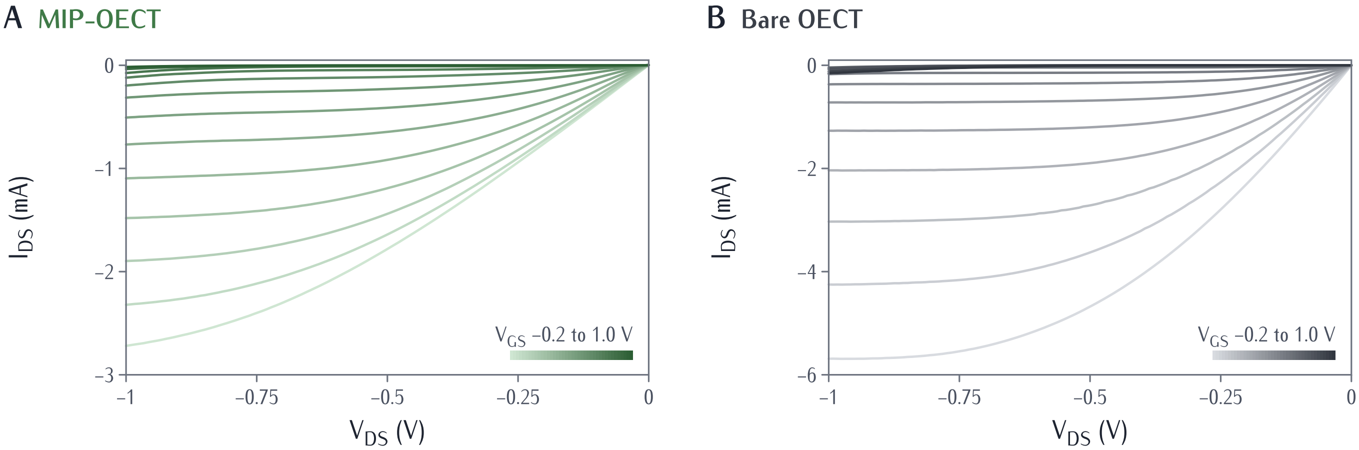


**Figure S2. Output characteristics of the OECTs.** $I_{DS}$ versus $V_{DS}$ for $V_{GS}$ from —0.2 to 1.0 V in 0.1-V steps (light to dark), in artificial sweat with 100 nM cortisol (SMU). (**A**) MIP-coated OECT. (**B**) Bare OECT.

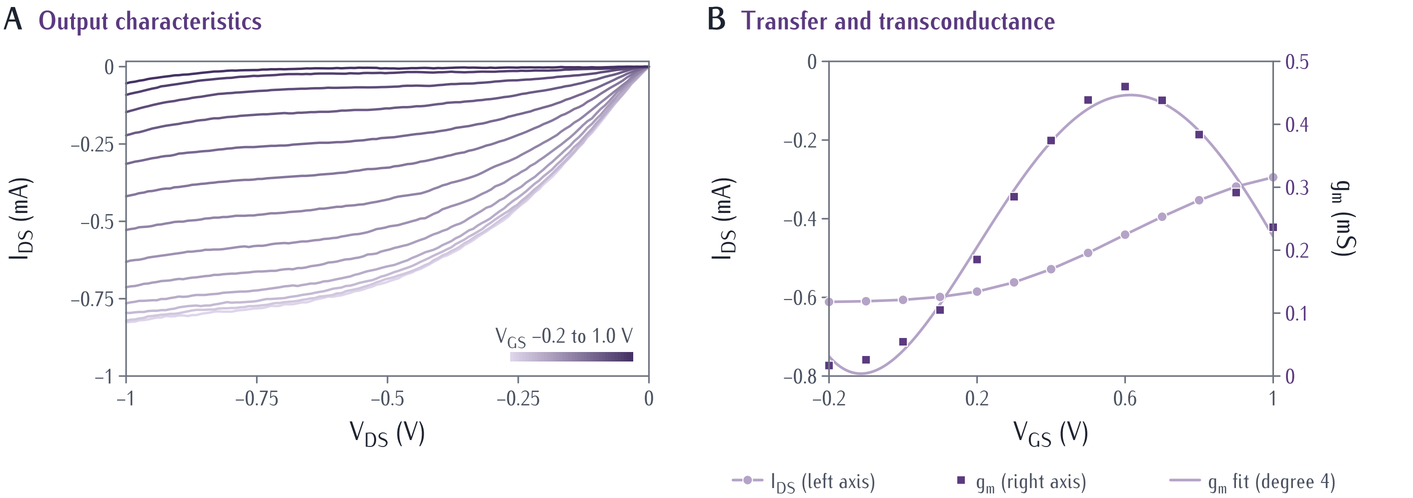


**Figure S3. Characteristics of the NIP-coated OECT.** (**A**) Output characteristics, as in Figure S2. (**B**) Transfer curve ($V_{DS} = -0.5$ V) and transconductance; line, fourth-order polynomial fit to $g_m$. Artificial sweat with 100 nM cortisol (SMU).

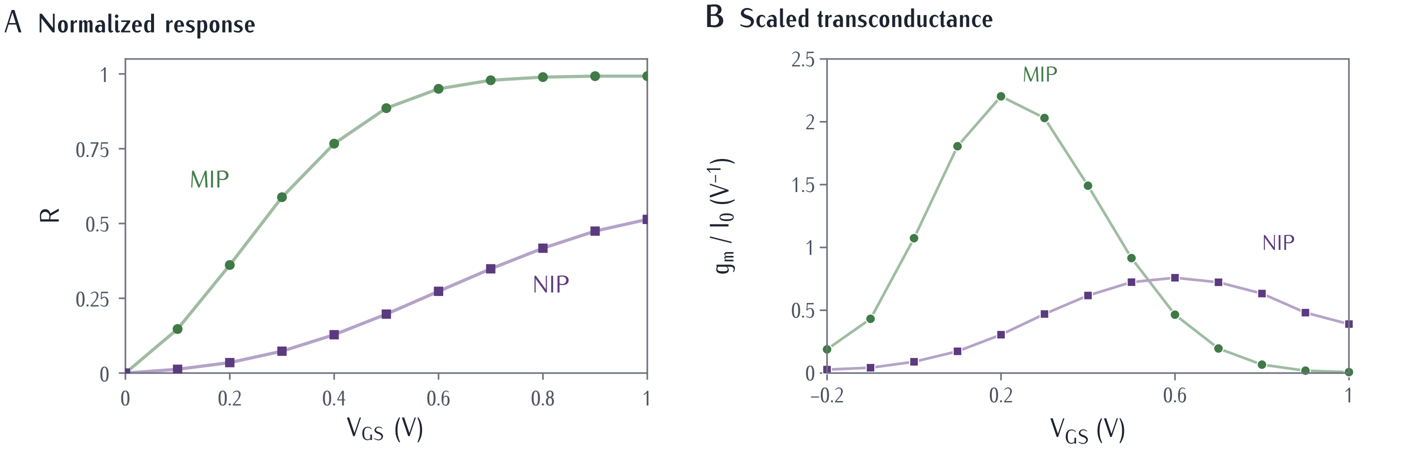


**Figure S4. Gating of the MIP- and NIP-coated OECTs.** (**A**) Normalized response $R$ versus $V_{GS}$ ($V_{DS} = -0.5$ V). (**B**) Transconductance normalized to $I_0$. The gate was stepped every 13 s for both devices.

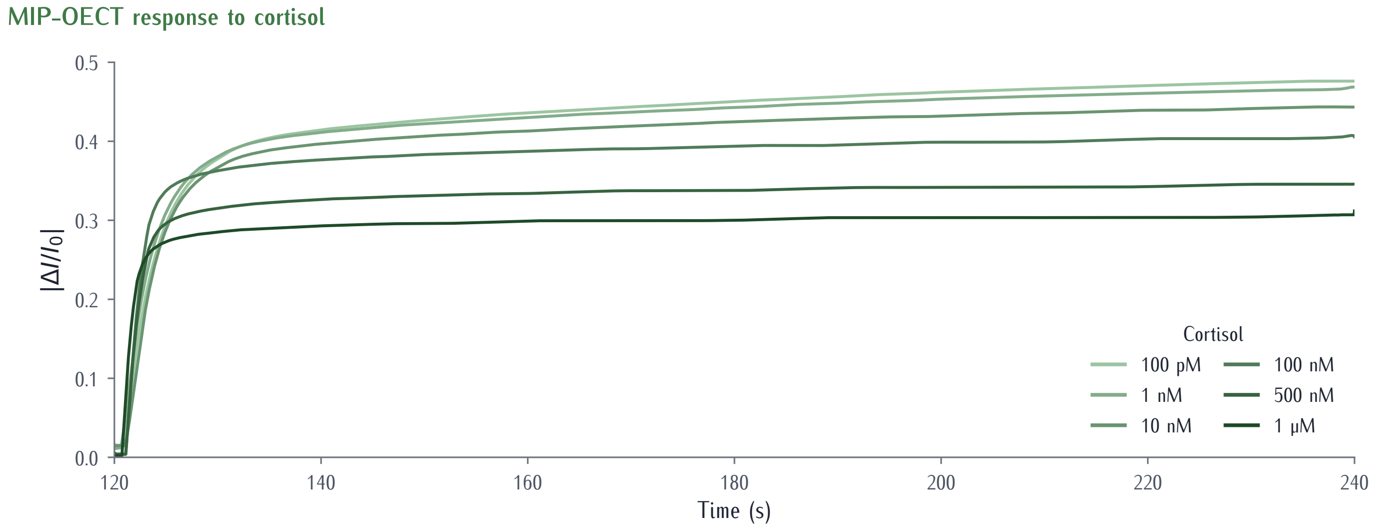


**Figure S5. Time response of the MIP-OECT (SMU).** Normalized response during the $V_{GS} = 0.2$ V phase for 100 pM to 1 $\mu$M cortisol in artificial sweat ($V_{DS} = -0.5$ V). Single recordings from one device, smoothed with a moving average that widens to 10 s.

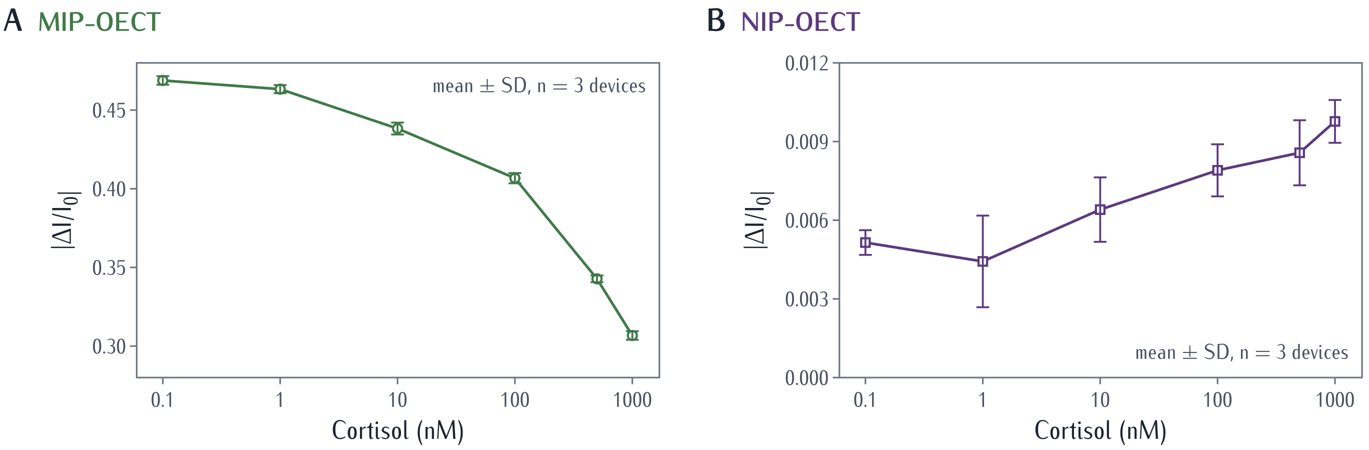


**Figure S6. Calibration of the MIP- and NIP-OECTs (SMU, day 1).** Normalized response versus cortisol concentration for (**A**) MIP-OECTs and (**B**) NIP-OECTs. Mean ± SD, $n = 3$ devices. Note the different vertical scales.

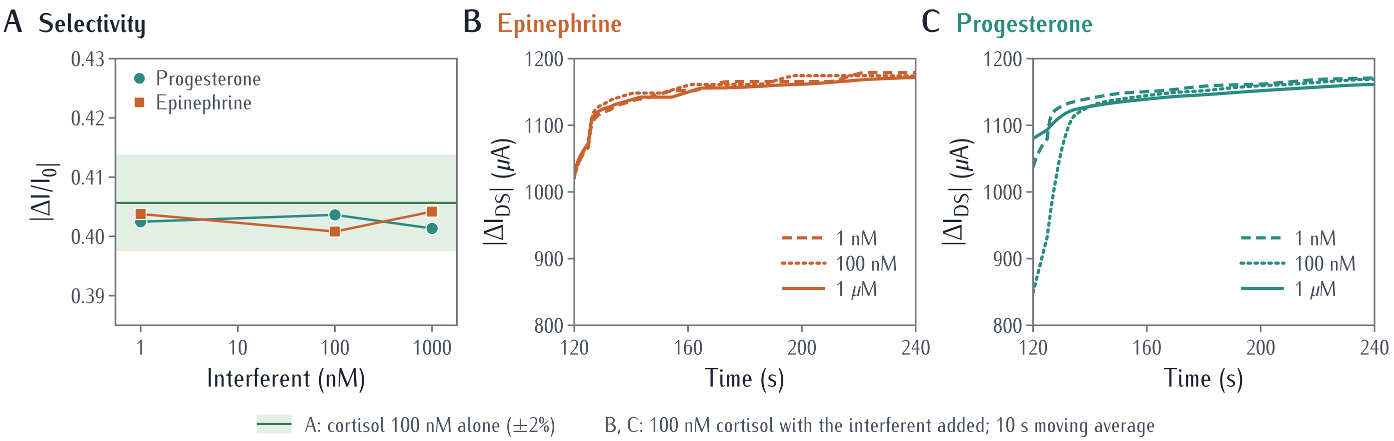


**Figure S7. Selectivity of the MIP-OECT.** (**A**) Normalized response to 100 nM cortisol in artificial sweat with epinephrine or progesterone added at 1 nM, 100 nM, or 1 $\mu$M. Solid line, 100 nM cortisol alone; shaded band, ±2%. (**B** and **C**) Drain-current change during the $V_{GS} = 0.2$ V phase with (**B**) epinephrine and (**C**) progesterone (10-s moving average).

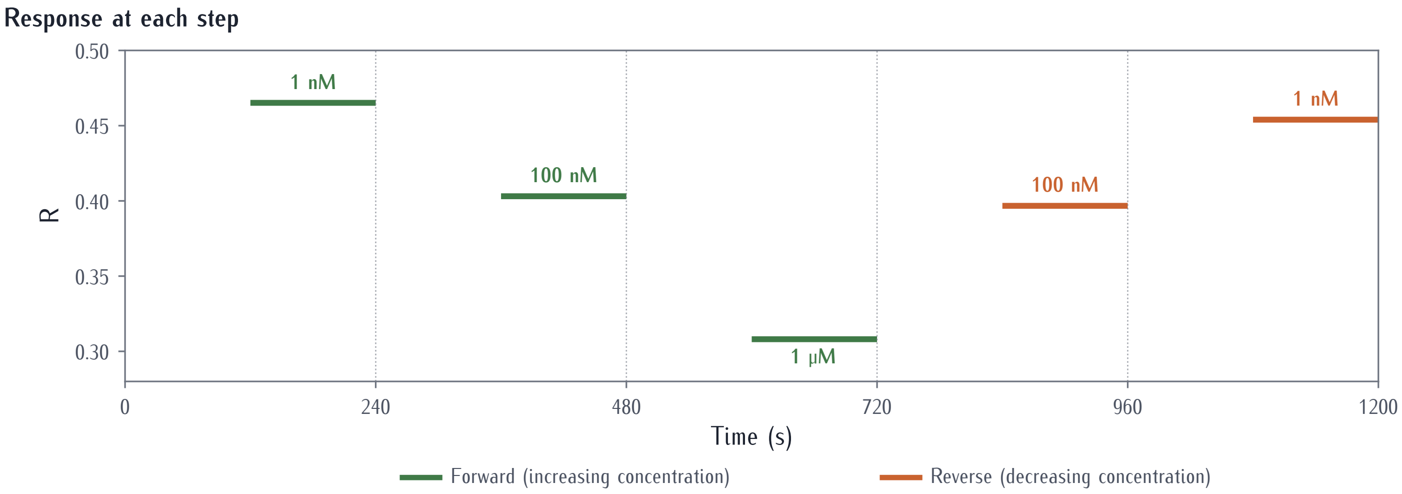


**Figure S8. Reversibility of the MIP-OECT response (SMU).** Response $R$ as cortisol was raised from 1 nM to 1 $\mu$M by adding cortisol to the drop on the device (forward, green) and lowered back to 1 nM by dilution with artificial sweat (reverse, orange). Each step comprised 120 s at $V_{GS} = 0$ V and 120 s at 0.2 V; values are plotted at the 0.2-V phase. Dotted lines, step boundaries.

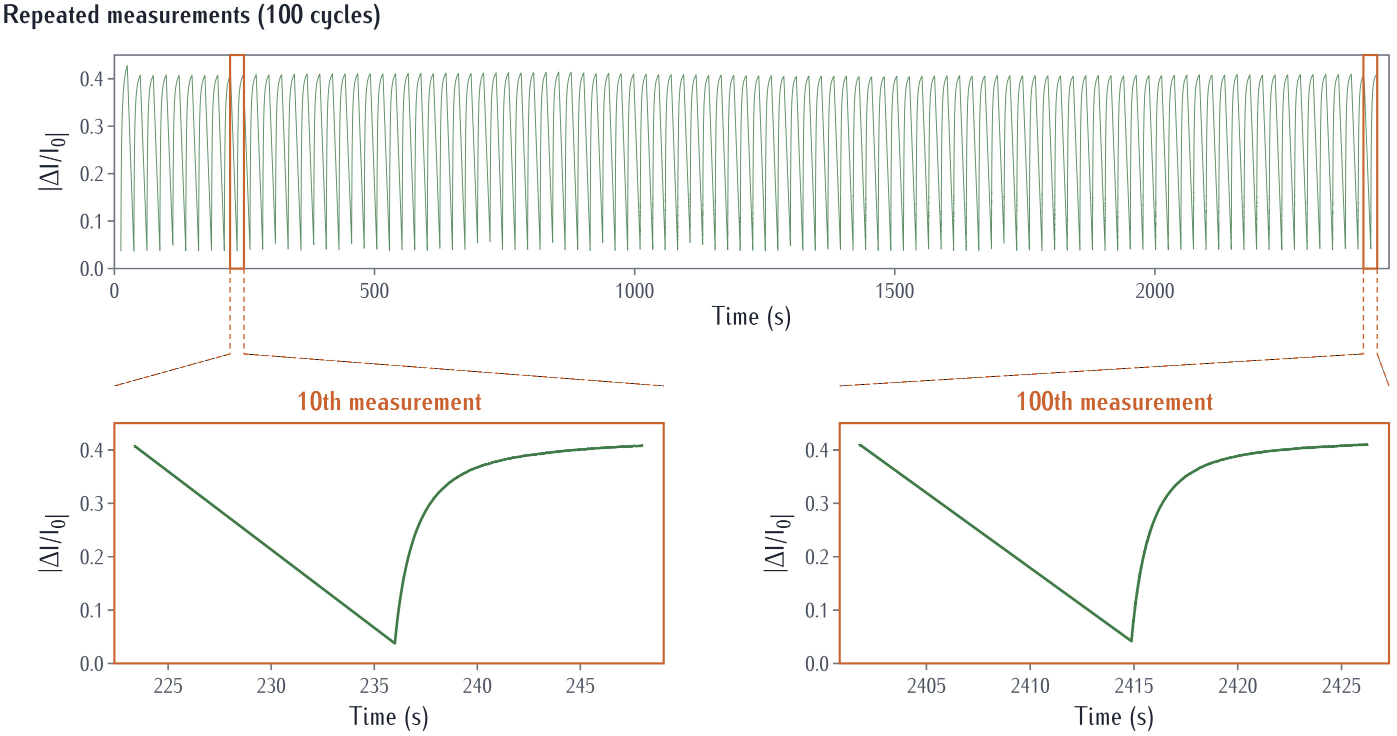


**Figure S9. Stability over 100 consecutive measurements (SMU).** Top, normalized response over 100 cycles in artificial sweat with 100 nM cortisol (0.2-V phase of about 12 s). Bottom, the 10th and 100th cycles (orange boxes, top). Straight segments join consecutive cycles across the 0-V phase, which is not plotted.

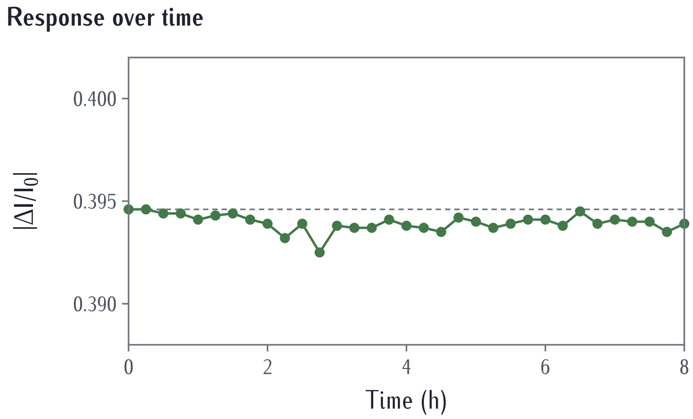


**Figure S10. Stability of the MIP-OECT over 8 h (wearable readout).** Normalized response measured every 15 min (33 readings) in artificial sweat with 100 nM cortisol. Dashed line, first reading.

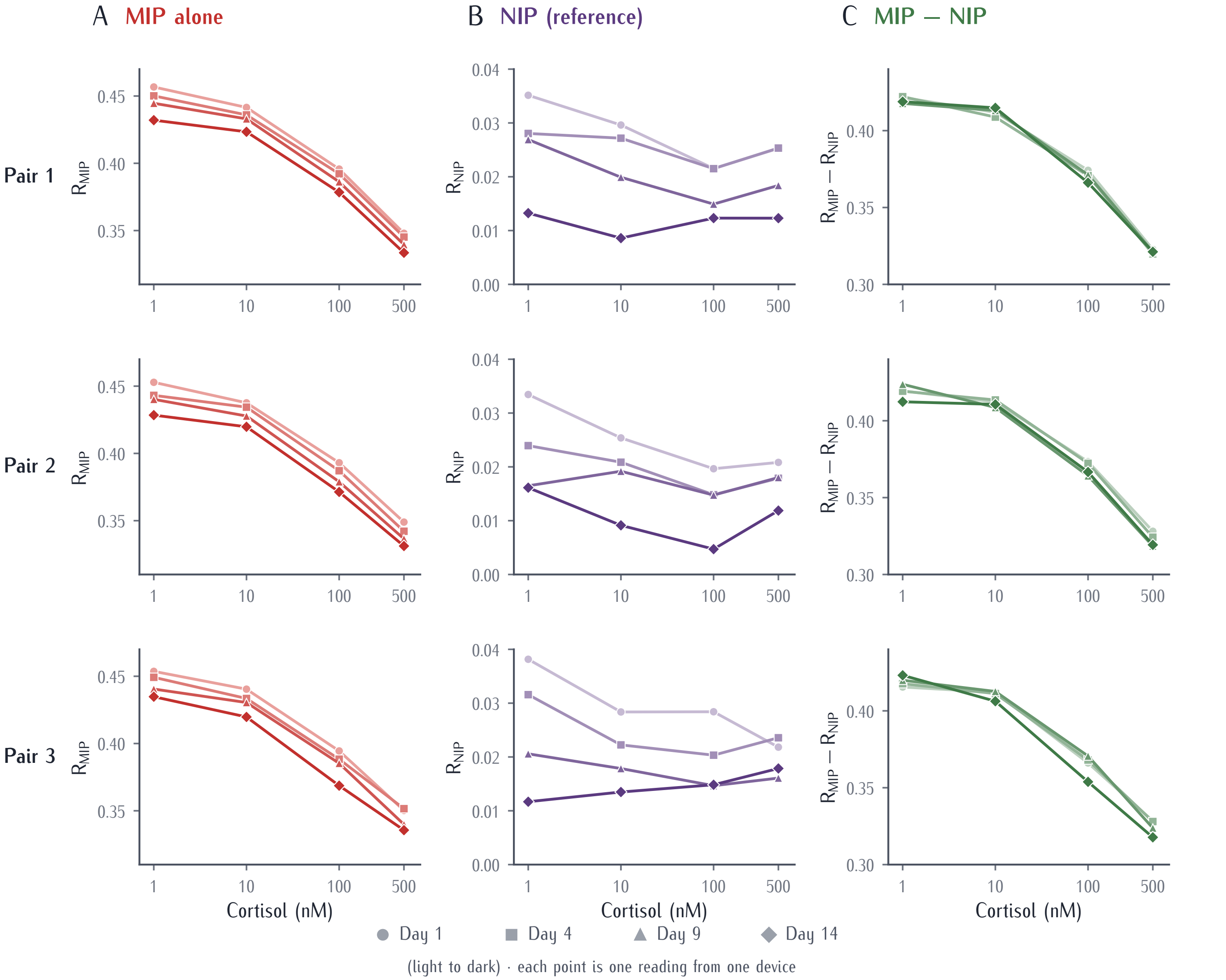


**Figure S11. Calibration of each device pair over two weeks.** (A) MIP, (B) NIP, and (C) MIP − NIP of each pair (rows) on days 1, 4, 9, and 14 (light to dark). Single readings; the means are shown in Figure 3B–3D.

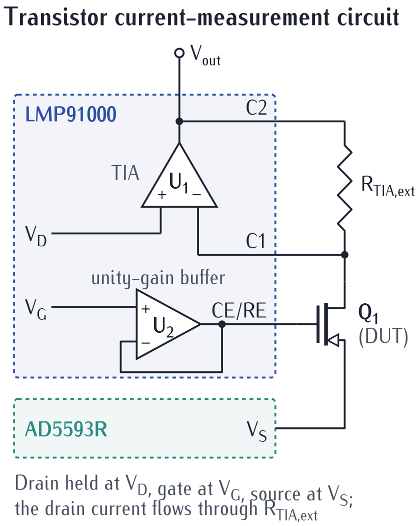


**Figure S12. Transistor current-measurement circuit.** Designators as in Note S2. The LMP91000 (dashed box) provides the transimpedance amplifier $U_1$ and the gate buffer $U_2$, and the AD5593R sets $V_S$. The full readout circuit is shown in Figure 4B.

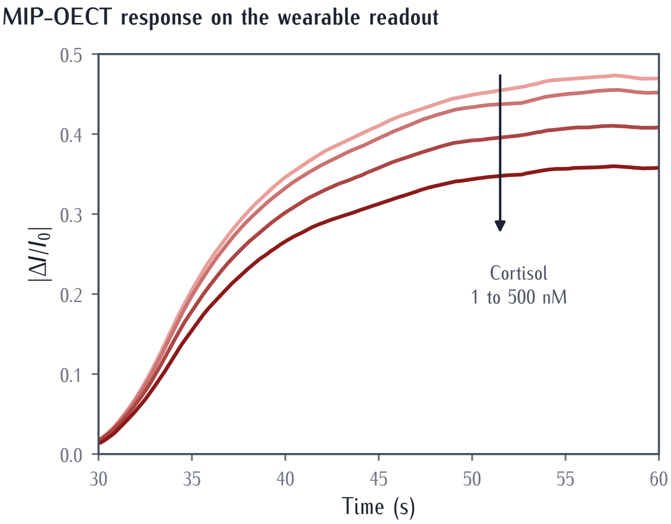


**Figure S13. Time response of the MIP-OECT on the wearable readout.** Normalized response during the $V_{GS} = 0.2$ V phase ($t = 30$ to 60 s) at 1, 10, 100, and 500 nM cortisol (light to dark) in artificial sweat; 5-s moving average; single readings from one device. Calibration values are the means of samples 1,483 to 1,800 (about 46 to 56 s; Experimental procedures).

**Table S1. Differential and internal-reference strategies in chemical sensors and biosensors, compared with this work.**

| Study | Sensor | Strategy | Reference element | Setting |
|---|---|---|---|---|
| *Transistor and electrochemical sensors* | | | | |
| Bergveld[1] | ISFET | Differential pair of a sensing ISFET and a reference FET (REFET); common signals such as an unstable reference-electrode potential cancel | Transistor with an ion-insensitive surface | Review of ISFET development; laboratory |
| Li et al.[2] | Electrochemical aptamer sensor | Dual-reporter correction: difference between a target-responsive and a target-insensitive redox reporter on the same sensor | Second redox reporter at a separate potential | Undiluted whole blood, *in vitro* |
| Arroyo-Currás et al.[3] | Electrochemical aptamer sensor | Kinetic differential measurement: difference between two frequencies that respond to the target in opposite directions but drift together | Same electrode, second interrogation frequency | *In vivo* (anesthetized and awake rats) |
| Li et al.[4] | Electrochemical aptamer sensor | Ratio of two interrogation frequencies, removing the need for per-sensor calibration | Same electrode, second interrogation frequency | Undiluted whole blood, *in vitro* |
| Zhao et al.[5] | Stretchable extended-gate OFETs (cortisol, glucose, $Na^{+}$) | Diode-connected transistor pair; difference between target and reference outputs suppresses drift from bias stress (20 min), temperature (25–40 °C), and strain | Transistor with a scrambled aptamer (cortisol) or no receptor | Wearable, sweat, on body (cold pressor) |
| *Imprinted-polymer sensors* | | | | |
| Seidler et al.[6] | MIP-coated QCM (yeast cells) | Imprinted and non-imprinted electrodes on one crystal, measured side by side; non-imprinted electrodes indicate non-specific binding | Non-imprinted electrode on the same crystal | Cell suspensions, benchtop |
| Sullivan et al.[7] | MIP nanoparticle SPR (antibiotics) | Signal of a reference flow channel subtracted from the imprinted channel | Channel prepared without imprinted nanoparticles | River water and milk, benchtop |
| Kang et al.[8] | MIP-coated photoelectrode (dopamine) | MIP minus weighted NIP photocurrent to cancel non-specific adsorption of interferents; also reported to remove baseline drift within a measurement | Separate NIP-coated photoelectrode | Serum samples, benchtop |
| Zhao et al.[9] | MIP-coated electrodes (two analytes) | Difference between two MIPs imprinted for different targets, each acting as the other's reference; baseline drift within a measurement also suppressed | Second MIP (no NIP) | Urine samples, benchtop |
| Yu et al.[10] | MIP-coated ECL electrode (dopamine) | MIP minus NIP after exponential transformation of the semi-logarithmic responses, to cancel interference | Separate NIP-coated electrode | Serum samples, benchtop |
| Parlak et al.[11] | MIP-coated OECT (cortisol) | NIP used as a separate control to confirm imprinting; not used for correction | Separate NIP device (control only) | Wearable, sweat, on body |
| **This work** | **Paired MIP- and NIP-coated OECTs (cortisol)** | **Continuous subtraction of the NIP response from the MIP response, tested against a single day-1 calibration over 14 days** | **Matched NIP-coated transistor** | **Artificial sweat (4 calibrations over 14 days); human sweat collected in 4 sessions over 14 days and read on a separate pair; one on-body session with the worn sensor** |

[2] Hui Li, Netzahualcóyotl Arroyo-Currás, Di Kang, Francesco Ricci, and Kevin W. Plaxco. Dual-reporter drift correction to enhance the

performance of electrochemical aptamer-based sensors in whole blood. *Journal of the American Chemical Society*, 138(49):15809–15812, 2016.

### Note S1. Drift metrics from the mean curves and from individual pairs

The drift metrics in the main text are calculated from the mean calibration curves: the three devices are first averaged on each day, and each metric is then calculated from these daily means. Here each metric is instead calculated for each device pair, pairing each MIP with its own NIP, and the result is averaged over the three pairs (Table S2; individual devices in Figure S11).

The two approaches give similar results. Per pair, the calibration range of the MIP alone decreased by $7 \pm 3\%$, whereas that of the differential output did not decrease on average ($-7 \pm 12\%$, that is, a mean increase, ranging from a 1% decrease to a 21% increase between pairs; mean ± SD, $n = 3$; paired $t$ test, $p = 0.13$), and the shift at 1 nM was $-21.5 \pm 2.8\%$ and $-0.6 \pm 8.5\%$ of the day-1 range ($p = 0.024$). The reduction in day-to-day spread per pair was $65 \pm 15\%$ at 1 nM ($p = 0.033$) and $70 \pm 3\%$ at 10 nM ($p = 0.002$), compared with 90% and 92% for the mean curves. The drift of the MIP alone had the same direction in all three devices, so averaging the devices does not reduce it, whereas the small day-to-day variation that remains in the differential output differs between pairs and partly cancels when the devices are averaged. The per-pair values therefore describe what a single sensor can be expected to achieve. The remaining spread of individual pairs, 0.004 to 0.011 at 1 and 10 nM, is about 11 to 28 times the readout noise of 0.0004 (Experimental procedures), so it reflects changes in the devices rather than measurement noise. At 100 and 500 nM the spread of individual pairs fell by $49 \pm 12\%$ ($p = 0.011$) and $53 \pm 24\%$ ($p = 0.045$).

**Table S2. Drift metrics for each device pair.** Sensitivity loss, decrease in the calibration range ($R$ at 1 nM minus $R$ at 500 nM) from day 1 to day 14, as a percentage of the day-1 range. 1 nM shift, change in the 1 nM response over the same period, as a percentage of the day-1 range. Spread, highest minus lowest of the four daily values. Reduction, decrease in spread from MIP to MIP − NIP. Negative values indicate an increase. $p$, two-sided paired $t$ test ($n = 3$ pairs).

| Metric | Output | Pair 1 | Pair 2 | Pair 3 | Mean ± SD | $p$ |
|---|---|---|---|---|---|---|
| Sensitivity loss (%) | MIP | 9.4 | 6.4 | 4.1 | 6.6 ± 2.7 | |
| | MIP − NIP | 1.3 | −1.9 | −21.0 | −7.2 ± 12.1 | 0.13 |
| 1 nM shift (% of day-1 range) | MIP | −22.69 | −23.48 | −18.23 | −21.47 ± 2.83 | |
| | MIP − NIP | −2.80 | −7.76 | +8.76 | −0.60 ± 8.47 | 0.024 |
| Spread at 1 nM | MIP | 0.0247 | 0.0244 | 0.0188 | 0.0226 ± 0.0033 | |
| | MIP − NIP | 0.0044 | 0.0114 | 0.0076 | 0.0078 ± 0.0035 | 0.033 |
| | Reduction (%) | 82.3 | 53.3 | 59.6 | 65.1 ± 15.3 | |
| Spread at 10 nM | MIP | 0.0181 | 0.0178 | 0.0207 | 0.0189 ± 0.0016 | |
| | MIP − NIP | 0.0060 | 0.0049 | 0.0064 | 0.0058 ± 0.0008 | 0.002 |
| | Reduction (%) | 66.8 | 72.5 | 69.2 | 69.5 ± 2.8 | |
| Spread at 100 nM | MIP | 0.0173 | 0.0217 | 0.0258 | 0.0216 ± 0.0043 | |
| | MIP − NIP | 0.0080 | 0.0093 | 0.0168 | 0.0114 ± 0.0047 | 0.011 |
| | Reduction (%) | 53.4 | 57.3 | 35.0 | 48.6 ± 11.9 | |
| Spread at 500 nM | MIP | 0.0145 | 0.0178 | 0.0160 | 0.0161 ± 0.0016 | |
| | MIP − NIP | 0.0029 | 0.0096 | 0.0107 | 0.0077 ± 0.0042 | 0.045 |
| | Reduction (%) | 80.2 | 46.1 | 33.0 | 53.1 ± 24.4 | |

**Table S3. Calibration slopes of each device pair.** Slope of $R$ versus $\log_{10}$ concentration (1 to 500 nM), per decade. Last rows, change from day 1 to day 14; $p$, two-sided one-sample $t$ test against zero ($n = 3$ pairs).

| Pair | Day | MIP | NIP | MIP − NIP |
|---|---|---|---|---|
| 1 | 1 | −0.0401 | −0.0043 | −0.0358 |
| 1 | 4 | −0.0387 | −0.0016 | −0.0370 |
| 1 | 9 | −0.0390 | −0.0036 | −0.0355 |
| 1 | 14 | −0.0367 | +0.0001 | −0.0367 |
| 2 | 1 | −0.0385 | −0.0049 | −0.0336 |
| 2 | 4 | −0.0378 | −0.0028 | −0.0350 |
| 2 | 9 | −0.0389 | 0.0000 | −0.0389 |
| 2 | 14 | −0.0368 | −0.0022 | −0.0346 |
| 3 | 1 | −0.0385 | −0.0053 | −0.0331 |
| 3 | 4 | −0.0367 | −0.0031 | −0.0336 |
| 3 | 9 | −0.0374 | −0.0019 | −0.0355 |
| 3 | 14 | −0.0379 | +0.0021 | −0.0401 |
| 1 | 1 → 14 | +0.0035 | +0.0044 | −0.0009 |
| 2 | 1 → 14 | +0.0017 | +0.0028 | −0.0010 |
| 3 | 1 → 14 | +0.0005 | +0.0075 | −0.0070 |
| Mean ± SD | | +0.0019 ± 0.0015 | +0.0049 ± 0.0024 | −0.0030 ± 0.0035 |
| $p$ | | 0.16 | 0.072 | 0.27 |

### Note S2. Transistor current-measurement circuit

A representative transistor current-measurement circuit is shown in Figure S12. Designators in this note refer to Figure S12 and differ from those in Figure 4B of the main text. The transistor $Q_1$ is the device under test (DUT). The source-bias voltage $V_S$ is generated using a DAC output; in this implementation, the AD5593R configurable ADC/DAC is used for this purpose. The gate-bias voltage $V_G$ is applied through the unity-gain buffer $U_2$. The drain-current readout is implemented using the transimpedance amplifier (TIA) $U_1$, with the desired drain-bias reference $V_D$ applied to its non-inverting input.

Under closed-loop operation, the negative feedback of $U_1$ maintains the inverting-input node $C1$ approximately at the reference potential $V_D$. Since the amplifier input current is negligible, the drain current from $Q_1$ flows predominantly through the external transimpedance resistor $R_{\mathrm{TIA,ext}}$, connected between the $C1$ and $C2$ terminals. The corresponding voltage difference at the TIA output is therefore proportional to the drain current, with

$$|V_{\mathrm{out}} - V_D| \approx |I_D|\, R_{\mathrm{TIA,ext}}, \tag{S1}$$

where the polarity of $V_{\mathrm{out}} - V_D$ depends on the direction of the drain current. In the implemented readout circuit, the potentiostat AFE LMP91000 provides the TIA and gate-drive functions represented by $U_1$ and $U_2$, respectively. Its CE and RE pins are connected together and drive the transistor gate, and its WE pin is connected to the transistor drain. Although the LMP91000 provides programmable internal transimpedance resistances from 2.75 kΩ to 350 kΩ, the required TIA resistance in this implementation is 1 kΩ, so an external 1 kΩ resistor $R_{\mathrm{TIA,ext}}$ is connected between its $C1$ and $C2$ pins. The LMP91000 is configured through the $I^2C$ interface, and its analog output $V_{\mathrm{out}}$ is sampled by the ADC of the SoC.
